# Codependence in the *Nephromyces* species swarm depends on heterospecific bacterial endosymbionts

**DOI:** 10.1101/2020.10.18.344572

**Authors:** Christopher Paight, Elizabeth Sage Hunter, Christopher E Lane

## Abstract

The phylum Apicomplexa encompasses 6000 ubiquitous animal parasites, including *Plasmodium*, the most deadly human parasite on Earth. Anciently parasitic lineages, like apicomplexans, lose core metabolic pathways over time, as they evolve less costly scavenging mechanisms. The recent description of a mutualistic apicomplexan, *Nephromyces*, from deep within this parasitic group, opened the possibility of an evolutionary innovation that allowed an escape from a parasitic lifestyle. Nuclear genome data from *Nephromyces*, as well as the three bacterial symbionts that live within this species complex, demonstrate that the bacteria within *Nephromyces* contribute essential cofactors and amino acids that have enabled *Nephromyces* to abandon a parasitic lifestyle. Among these, bacterial lipoic acid appears to be a key cofactor for the reduction of virulence in *Nephromyces*. However, whereas we use FISH microscopy to reveal that each individual *Nephromyces* harbors no more than one endosymbiont type, no single bacterial endosymbiont can account for all missing metabolites. Based on the unique habitat of *Nephromyces*, as well as genomic, culturing, and wild population data, we conclude that *Nephromyces* has evolved as an extraordinary clade of codependent species, unlike any previously described.

## Introduction

Symbiosis is one of the most important evolutionary processes shaping biodiversity on Earth. Symbiotic communities often bring together organisms from different domains of life, which can provide an unparalleled route to evolutionary innovation, allowing life to thrive in unconventional niches such as hydrothermal vents (1) or the oxic-anoxic interface (2). One such extreme environment is the renal sac of the marine invertebrate genus *Molgula*. These tunicates produce a liquid-filled organ, which contains high enough concentrations of uric and oxalic acids to form crystal structures analogous to human kidney stones (3). Surprisingly, within this environment live multiple apicomplexan species of *Nephromyces* (Fig. 1), with vertically transmitted bacterial endosymbionts living inside them (4). Unlike the ~6000 described species of apicomplexans, *Nephromyces* spp. are not parasitic (5). Instead, they maintain a mutualistic or commensal relationship with molgulid tunicates, using a combination of retained metabolic pathways and their bacterial endosymbionts (6).

**Figure 1:**
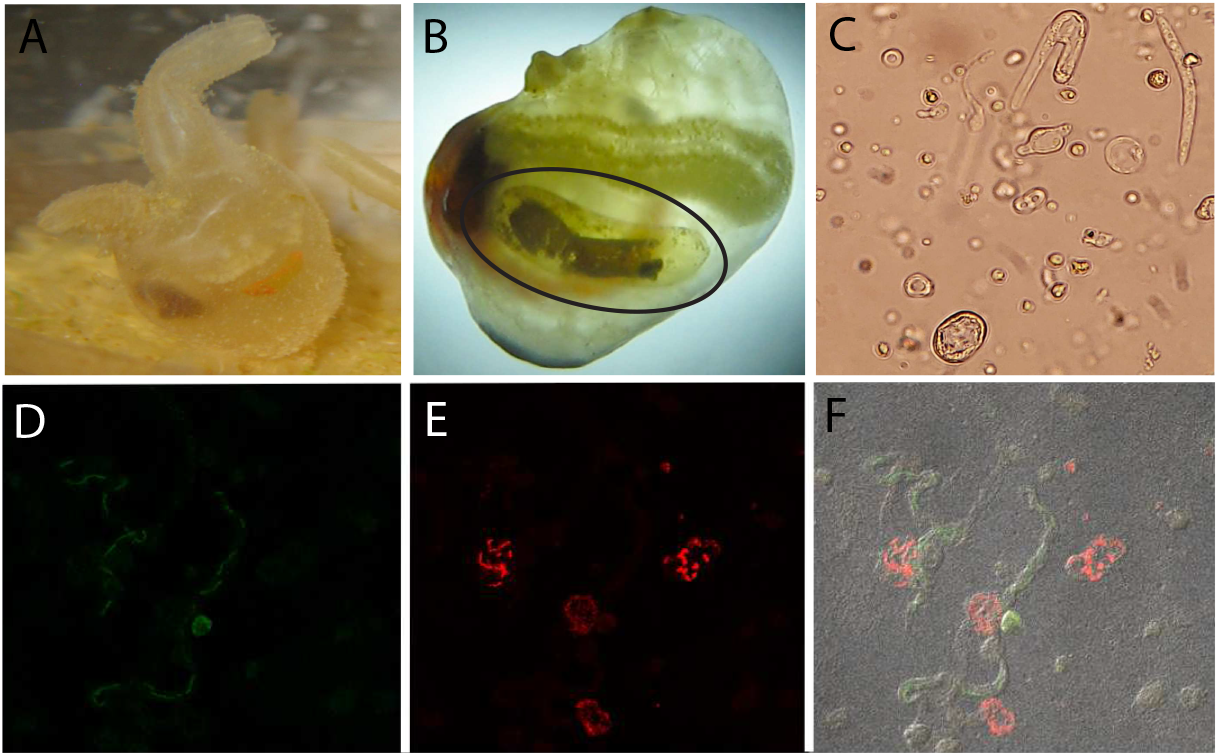
A) Laboratory cultured *Molgula manhattensis*, one of the *Mogula* tunicate hosts examined in this study. B) *Molgula manhattensis* with tunic removed to show renal sac (circled), where *Nephromyces* spp. spend their entire life cycle. The brown substance inside the renals sac are concretions of crystallized uric acid and calcium oxalate. C) Light microscopy photo of several *Nephromyces* life stages, all cells in photo are of *Nephromyces*. Bacteiodetes (D) and alphaprotebacterial (E) endosymbionts labeled with 16S rRNA class specific probes, using fluorescence *in situ* hybridization. F) The composite image of (D) and (E) showing these endosymbionts localized to different *Nephromyces* cells.

Besides *Nephromyces*, the phylum Apicomplexa is composed entirely of obligate metazoan parasites. As a result of an estimated 800 million years of evolution as obligate parasites, many of the genomic patterns associated with the response to a parasitic lifestyle have been described from the genomes of apicomplexan species (7). Apicomplexan-specific examples include expansions in the *Plasmodium* var protein family, which are involved in host manipulation and evasion, as well as an expansion of rhoptry, microneme, and dense granule proteins (8, 9). The list of core biosynthetic pathways apicomplexans have lost includes purine biosynthesis, purine degradation, biosynthesis of many amino acids, and vitamin biosynthesis (10–13). These losses make the parasite dependent on the host, not only for primary carbon and nitrogen, but also for any metabolites it can no longer generate, either by *de novo* synthesis or conversion. High demand on the host for these metabolites to fuel parasite growth increases the cost of infection, thereby increasing virulence. Parasites must maintain a delicate balance between transmission, virulence, and host immune system evasion.

The trade-offs in this balance have been described in detail (14–16), but one common solution many parasites adopt is maintaining low relative abundance inside the host. Higher parasite abundance will increase the cost to that host, and increase virulence. If the parasites kill the host before completing their lifecycle or before transmission to a new host, their fitness falls to zero. Similarly, if parasites have a high prevalence in a population and high lethality, they risk decimating their host population. High-sustained infection prevalence is a good indicator of low virulence (17), and low virulence is often achieved by self-limited reproduction by the parasites. In this way, *Nephromyces* stands out as a very atypical parasite.

*Nephromyces* has a nearly 100% infection rate, which is sustained almost year-round (5). Based on typical host/parasite dynamics, *Nephromyces* also reaches unexpectedly high cell densities. These atypical epidemiological factors were the basis for the conclusion that *Nephromyces* must be mutualistic (5). In order to reach high cell densities while maintaining low virulence, *Nephromyces* was predicted to produce something of high value to the host, to offset the cost associated with maintaining such high densities of an obligate parasite. *Nephromyces* is also unusual because it lives the majority of its life extracellularly (18). Whereas some early-diverging gregarines are primarily extracellular, the Nephromycidae resolves as sister to the Hematozoans, a clade of obligate intracellular blood parasites (19). While it remains unknown if *Nephromyces* is capable of forming direct connections to tunicate host cells, there is no microscopic support for direct cell-to-cell connections. Instead, *Nephromyces* cells are found independent of tunicate cells in the lumen of the renal sac. The renal sac is a large ductless structure of unknown function exclusively found in a Molgulidae family of tunicates (20).

Despite its name, the renal sac does not function as a typical renal organ but was named for the large deposits of crystallized uric acid and calcium oxalate. Many ascidians have localized deposits of uric acid, but tunicates in the Molgulidae family have the largest uric acid deposits (21). While the function of these deposits of uric acid in the tunicate remain unclear, previous work demonstrated that *Nephromyces* is able to degrade uric acid, because it retains the ancestral purine degradation genes lost in all other apicomplexans (6). Based on transcriptome data and pathway analysis, uric acid may be the primary source of carbon and nitrogen for *Nephromyces* (6). Uric acid is an atypical source of carbon and nitrogen, but not completely unknown. There are several species of bacteria and fungi that can be cultured on media only uric acid (22–24).

The shift from an intracellular environment rich in preformed metabolites and precursor molecules, to an extracellular lifestyle inside the renal sac, likely limited the amount and type of preformed metabolites available for the ancestor of *Nephromyces* to scavenge. Given the large number of basic biosynthetic pathways lost in apicomplexans, this organism would have needed to evolve mechanisms of obtaining these essential metabolites not encoded in its genome. Metabolites could still be scavenged from the host; while not as abundant, many essential metabolites are still present in the extracellular environment (25). Another potential source of these metabolites could be the development of *de novo* synthesis pathways. However, the sheer number of essential pathways missing in apicomplexans makes the transition to extracellular unlikely. A third option for filling the need for so many basic metabolites is the acquisition of bacterial endosymbionts.

Preliminary fluorescence *in situ* hybridization microscopy performed on *Nephromyces* indicated species harbor three bacterial endosymbionts: an alphaproteobacteria endosymbiont (N *α* e), Betaproteobacteria (N *β* e), and a Bacteroidetes (Nbe). Acquisition and maintenance of bacterial endosymbionts is a common way for eukaryotes to gain new metabolic pathways and capabilities. The functional capabilities of bacterial endosymbionts exploited by eukaryotic hosts include amino acid metabolism and vitamin metabolism (26), nitrogen metabolism (27), defense (28), chemotrophic energy production (29), and photosynthesis (30), to name a few. Whereas bacterial endosymbionts are common in many protist lineages, they are unknown in the phylum Apicomplexa outside the Nephromycidae. Previous speculation (5) that *Nephromyces* bacterial endosymbionts are responsible for the high levels of purine degradation observed have recently been rejected (6). However, the bacterial endosymbionts were likely instrumental in the transition from intracellular to extracellular and colonization of the renal sac.

Here we show that the diversity of *Nephromyces* spp. and their endosymbionts are unexpectedly high within each renal sac and forms a codependent community within their host. We determined that as many as 20 different species of *Nephromyces* inhabit a typical renal sac. Whereas *Nephromyces* species can host three different types of bacterial endosymbionts, FISH microscopy allowed us to show that only a single bacterial type exists within any one individual. Through the reconstruction and analyses of the endosymbiont bacterial genomes, we infer that each type supplies its *Nephromyces* host with different metabolites.

Given that no individual species of *Nephromyces*, in combination with its endosymbiont, can produce a complete set of essential amino acids, we hypothesize that each of them depends on multiple congeners to meet its needs. Growth experiments demonstrate that individual *Nephromyces* species cannot form a viable infection. *Nephromyces* has thus evolved as an extraordinary clade of codependent species.

## Results

### F*IS*H Microscopy

Fluorescence *in situ* hybridization (F*is*H) of wild *Molgula manhattensis* tunicates revealed persistent co-infections of alphaproteobacteria and Bacteroides. While these two classes of bacterial endosymbionts were frequently found together in the same renal sac, they never co-occurred in the same *Nephromyces* cell. A betaproteobacteria probe was also used in all experiments, but was never visualized in our samples, in contrast to (31). Based on the corresponding amplicon data for tunicates, we believe this was a problem with probe specificity and binding, and not a genuine absence of betaproteobacteria endosymbionts. All three probes have successfully hybridized in previous experiments (31), and no individual *Nephromyces* cells were found to contain more than one class of endosymbiont.

### RNA Sequencing Results

High-quality RNA was extracted from the contents of a single renal sac, resulting in 195,694 transcripts from *Molgula manhattensis, Nephromyces*, and the bacterial endosymbionts. After binning by species, 60,223 transcripts were attributed to *Nephromyces*, and 6,589 were attributed to the bacterial endosymbionts. The large number of transcripts attributed to *Nephromyces* was due to multiple species infecting a single host (estimated 69% duplication). Clustering by percent sequence identity resulted in 26,938 at 90%, 23,850 at 80%, 21,762 at 70%, 19,540 at 60%, 16,668 at 50%. Due to the multi-species community, the transcriptome is from several species of *Nephromyces*, but we estimate that there are between 8,000 and 12,000 unique transcripts in *Nephromyces*. KEGG functionally predicts 6,987. BUSCO was used to assess the completeness of *Nephromyces* transcriptome resulting in 81.8% complete transcripts and 6.3% partial.

### *Nephromyces* Genome

Due to the multispecies nature of the system, the genome and transcriptome for *Nephromyces* are pan-genomic and pan-transcriptomic, and we cannot rule out that some of our assembled contigs and transcripts are potentially chimeric. However, these limitations do not limit our characterization of the metabolic contribution of *Nephromyces* as a genus. The *Nephromyces* pan-genome assembled into 2,156 contigs has an N50 50,747 bp with 36% GC content, a sum length of 60,134,441 bp, and BUSCO estimated completeness of 51.1% and duplication is 13.5%.

There are 3,070 predicted protein-coding genes in the *Nephromyces* dataset. Of these protein-coding genes, 2,834 were recovered in the transcriptome, 236 are unique to the genome, and 794 protein-coding genes in the transcriptome are missing from the genome. The incompleteness of the genome makes determining the true absence of genes impossible, but by combining it with the more complete transcriptome, we are able to better predict the metabolic capabilities and contributions of *Nephromyces* to the system. The genome also provides support that transcripts binned as *Nephromyces* based on phylogeny were binned correctly.

### Endosymbiont Genome

The genomes of three bacterial endosymbionts were retrieved from the metagenome of *Nephromyces*. These genomes were small, gene dense, and AT-rich. The presence of two closely related *α*-proteobacteria with different genome organization limited our ability to assemble the genome onto a single contig. The *α*-proteobacteria bacteria assembled into 11 contigs ranging from 13kb to 312kb, with an estimated 995,540 bp genome size, 844 predicted protein-coding genes, and 25% GC content. The Nbe genome is circular, 495,352 bp in length, contains 503 protein-coding genes, and has a 22% GC content. The N *β* e genome is circular, 866,396 bp in length, contains 880 protein-coding genes, and has a 30% GC content. Each genome contained hypothetical proteins or uncharacterized proteins making up 14-46% of predicted protein-coding genes (S Table 1).

### Amplicon Diversity

Amplicon sequencing of Run One resulted in 25,895,690 reads with an average reads per sample of 137,743. After binning, there were 4,930,010 CO1 reads, 6,491,918 18S reads, and 14,468,228 16S reads. Following assembly in dada2 and decontamination, there were 1,876,107 sequences corresponding to 329 amplicon sequence variants (ASVs) for 18S, 1,522,378 sequences corresponding to 188 ASVs for CO1, and 62,905 sequences with 152 ASVs for 16S. The 18S data indicated that *Nephromyces*, like *Plasmodium* (32), encodes multiple distinct copies of 18S rRNA, making this marker uninformative for species quantification. This maker was subsequently dropped for Run Two and the data are not presented here.

A total of 335 distinct *Nephromyces* COI ASVs were recovered from samples collected from *Molgula manhattensis*. Before clustering there was an average of 79.26 ASVs, with a max of 143 ASVs and min of 27 ASVs, per tunicate host (Fig. 3). Clustering at the 98% and 97% identity level resulted in an average of 5.1 and 2.8 ASVs per tunicate (Fig. 3). The most common AVSs were found in 53% of tunicates sampled and the rarest in 2.12%. At the 97% level, the most common clusters were found in 75.5% of tunicates the rarest in 2.12%. We identified a total of 152 16s sequences from the bacterial endosymbionts: 49 from Betaproteobacteria, 89 from Alphaproteobacteria, and 14 from Bacteroidetes. The average number of 16S sequences per tunicate was 13.94 with a max of 49 and a min of 0 (Fig. 3). Forty percent of the tunicates sampled contained all three types of bacterial endosymbiont, and 90% contained at least two of the bacterial endosymbiont types. Out of the 50 *M. manhattensis* sampled for amplicons, only two did not contain any 16s endosymbiont sequences, despite containing *Nephromyces* CO1 amplicons.

The 52 *Molgula occidentalis* samples combined from Runs One and Two had a total of 182 distinct COI ASVs from *Nephromyces*. Before clustering there was an average of 26.79 ASVs, with a max of 82 and min of 8, per tunicate host (Fig. 3). When clustered to the 97% identity level there are a total of 19 clusters with an average of 7.8 per tunicate (Fig. 3). The most common AVSs were found in 75% of tunicates sampled and the rarest ASVs were in 1.9%. After clustering at 97%, the most common clusters were in 100% of tunicates the rarest in 1.9%. From a total of 64 16s rRNA sequences classified as bacterial endosymbionts, 24 resolved as Betaproteobacteria, 37 as Alphaproteobacteria, and 13 as Bacteroidetes. The average number of 16S sequences per tunicate was 16.7 with a max of 30 and a min of 3 (Fig. 3). Nearly 41% of samples contained all three types of bacterial endosymbiont, and 80.77% contained at least two of the endosymbiont types. All *M. occidentalis* collected had at least one bacterial endosymbiont 16S sequence.

### *Molgula manhattensis* laboratory culturing

Regardless of infection status all cultures of *M. manhattensis* reproduced successfully with no apparent differences. Culture with the full *Nephromyces* inoculum never lost *Nephromyces* (in one case over 4.5 years, ~9 generations). An average of 25ng of DNA was extracted from the renal fluid of these tunicates. Cloning *Nephromyces* COI sequence and the bacterial 16S found this culture had 4 different *Nephromyces* species two with alphaproteobacteria endosymbionts and two with Bacteroidetes. On the plate inoculated with the 1/100 dilution *Nephromyces* transmission was also stable and never lost (4.5 years, ~9 generations). DNA extractions of renal fluid from these animals was never detectable by qbit and cloning revealed two *Nephromyces* species one with alphaproteobacteria and one with betaproteobacteria. Only two cultures were evey successfully inoculated with a single *Nephromyces* species and *Nephromyces* was lost from both after two to three generations.. The single-species cell densities were so low that infection status had to be confirmed with PCR amplification as visual inspection with a dissecting microscope often missed the presence of *Nephromyces*. In both cases the bacterial endosymbiont could not be determined and may not have actually contained a bacterial endosymbiont.

## Discussion

Our combined genomic, microscopy and population data reveals a unique ecosystem within the renal sac of molgulid tunicates, where multiple species of *Nephromyces*, harboring one of three different strains of bacterial endosymbiont, work in concert. The genomes of the three bacterial endosymbiont types are highly reduced, but perform several key functions, as well as complete pathways partially encoded by *Nephromyces* (Fig. 2). Only N β eis capable of synthesizing a full complement of amino acids (with the exception of tryptophan) with *Nephromyces*, but FISH microscopy shows only a single endosymbiont type exists within each individual (Fig. 1). Additionally, N β e is missing genes to synthesize essential cofactors, such as lipoic acid, encoded by Nbe and N`a`e. Together, these data indicate that, not only do individual *Nephromyces* exchange metabolites with their endosymbiotic bacteria, but between conspecifics within the renal sac.

**Figure 2:**
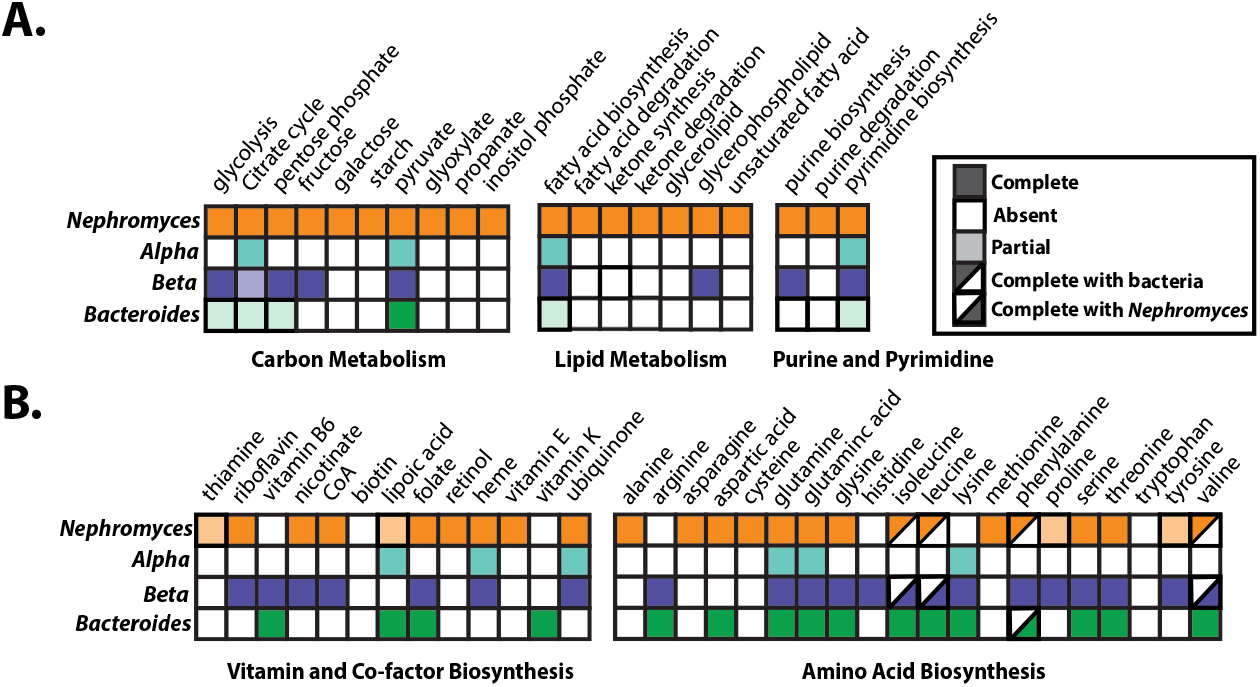
Graphical representation of metabolic pathways in *Nephromyces* and the three types of bacterial endosymbionts. Filled boxes represent complete pathways, empty boxes indicate no genes in the pathway are encoded, light shading represent partially complete pathways, and half filled boxes indicate pathways that require genes encoded by *Nephromyces* and a bacterial endosymbiont. A) Carbon, lipid and nucleotide metabolism showing greatly reduced capabilities of the bacterial endosymbionts. B) Amino acid, vitamin, and cofactor metabolism showing strong complementary biosynthetic integration of the bacterial endosymbionts and *Nephromyces*.

In support of the importance of multiple species infecting each renal sac, wild tunicates are either infected by multiple species of *Nephromyces* or have not yet been infected (Fig. 3). In experiments with laboratory raised tunicates, limiting the number of *Nephromyces* species per host affected cell densities and transmission efficiency. Single-species *Nephromyces* inoculations in lab raised tunicates grew poorly and were lost in subsequent generations of tunicates. Limited infections with two *Nephromyces* species were transmitted to other tunicates, but contained an order of magnitude fewer cells (measured by DNA quantities from extractions) than tunicates colonized by four or more species with undiluted renal fluid from wild tunicates.

**Figure 3:**
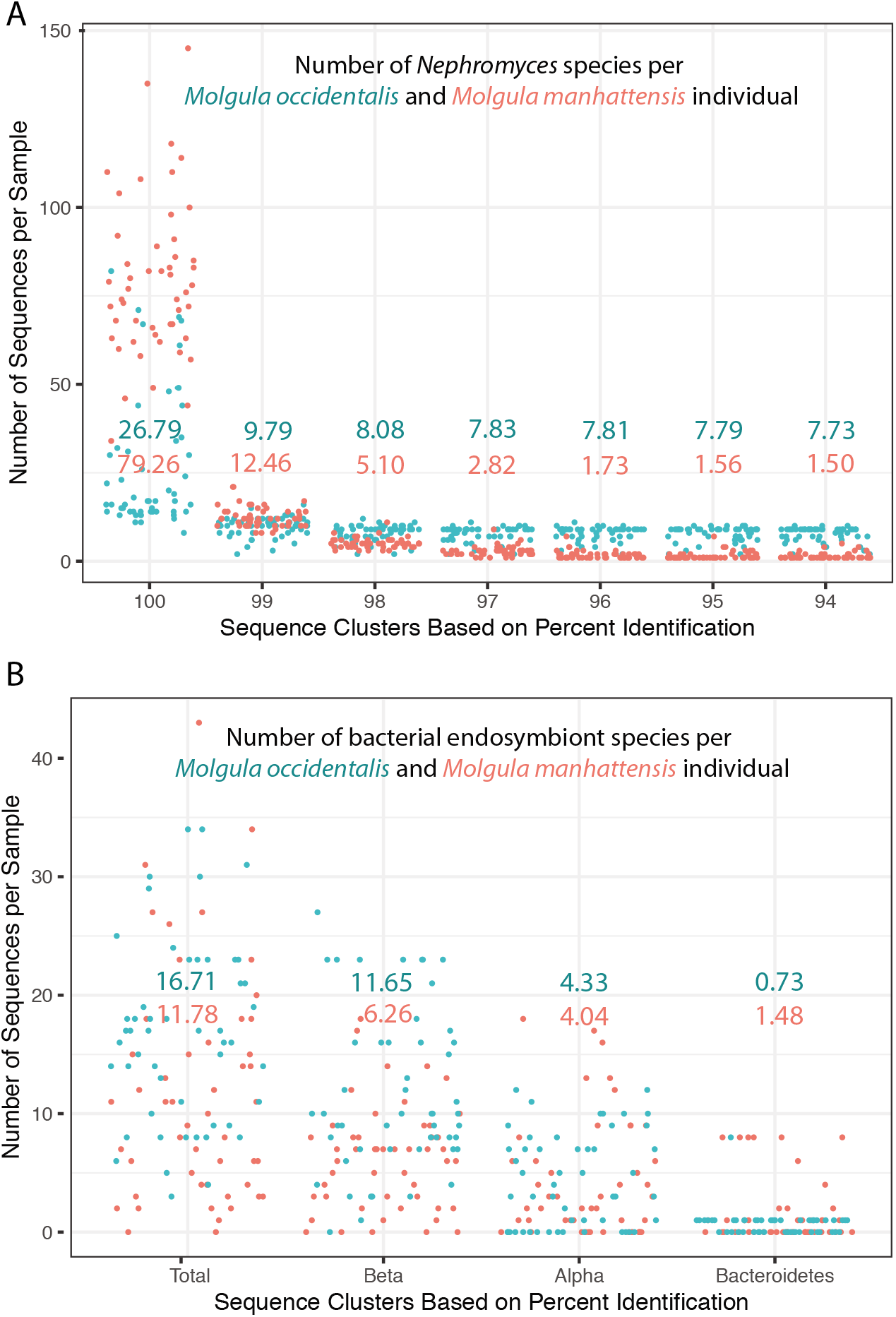
Biodiversity data across *Nephromyces* and their bacterial endosymbionts. A) Number of *Nephromyces* ASVs per tunicate individual (blue = *M. occidentalis*, red = *M. manhattensis*). The x-axis shows the different percent identity levels that the ASVs were clustered at. The mean at every clustering level for each host species is shown. B) The number and type of bacterial endosymbiont found from each tunicate host. Bacterial ASV’s were not clustered by percent identity. The mean for each host is shown.

The multi-species communities *Nephromyces* forms are a challenge for genomic methods. *Nephromyces* cells have highly variable morphology (Fig. 1) and rupture rapidly when manually isolated from renal sac fluid. Whereas the transcriptomic data reported here are from a single renal sac, both the genomic and transcriptomic sequences represent a community of closely related *Nephromyces*. By sequencing multiple species we have recovered the genomes of all three of the bacterial endosymbionts of *Nephromyces*, providing key insights into how this system functions *in vivo*.

The estimated 8,000 gene transcriptome for *Nephromyces* is largely complete as estimated by BUSCO. This number is similar to some of the most gene-rich apicomplexans, such as *Toxoplasma gondii*, which also encodes 8,000 genes. This is interesting given the phylogenetic placement of *Nephromyces* in the Hematozoa (19). Hematozoa, which contains the plasmodiidae and piroplasmida lineages, have some of the smallest genomes with the least number of genes of any sequenced apicomplexans. High gene numbers may be reflective of the greater biosynthesis and metabolic capabilities needed for an extracellular lifestyle and horizontal infection via spores in the water column.

Two of the biggest surprises in these datasets both involve purine metabolism. *Nephromyces* have the metabolic capabilities to convert xanthine into glyoxylate. Glyoxylate can be converted with serine-pyruvate aminotransferase (AGXT) into glycine and pyruvate, and *Nephromyces* additionally encodes malate synthase (MLS), which combines Glyoxylate and acetyl-CoA into malate. This is proposed to be the primary route of carbon, nitrogen, and energy acquisition for *Nephromyces* (6). This pathway is absent in all available apicomplexans genomes, but it appears the enzymes in this pathway were retained from the last common ancestor of Apicomplexa and not the result of a more recent horizontal gene transfer (6). The second surprise in *Nephromyces* purine metabolism is *de novo* purine biosynthesis (Sup Fig 1). While some apicomplexan lineages have one or two genes to synthesize inosine monophosphate (IMP) from immediate precursors, *Nephromyces* is predicted to encode the entire *de novo* purine synthesis pathway from 5-Phosphoribosyl diphosphate (PRPP). As the inability to synthesize purines has been widely targeted for drug development against other apicomplexan species, its presence in *Nephromyces* is unexpected.

The presence of both purine degradation and purine synthesis could be critical to the unusual epidemiology of *Nephromyces*. Tunicates lack the enzymatic ability to degrade purines past uric acid. By obtaining the bulk of the required carbon, nitrogen, and energy from a tunicate metabolic waste product, *Nephromyces* is able to limit impact on its host, while still reaching high cellular densities. *De novo* synthesis of purines means *Nephromyces* is not dependent on the host for IMP. Purine degradation and purine biosynthesis may have been the critical factors which allowed *Nephromyces* to leave the intracellular environment and colonize the renal sac.

Another critical factor in the ability of *Nephromyces* to survive in the renal sac is likely its bacterial endosymbionts. The N *α* e found in *Nephromyces* and *Cardiosporidium* are monophyletic, indicating that they have been maintained and vertically transmitted since the divergence of *Nephromyces* and *Cardiosporidium* (33). In addition to the N*α*e, *Nephromyces* has also acquired a N*β*e and a Nbe endosymbiont. The N*α*e and Nbe show a marked reduction in carbon metabolism with N α e only encoding genes for the citric acid cycle, and Nbe only capable of processing three-carbon compounds and encoding a partial citric acid cycle (Fig. 2). Such pronounced reduction suggests that *Nephromyces* provide their symbionts a limited ‘diet’ (26). In both symbionts, carbon metabolism may be dependent on pyruvate, which is one of the end products from the degradation of uric acid. All three of the bacterial endosymbionts paradoxically encode complete fatty acid biosynthesis but lack fatty acid degradation (Fig. 2). Presumably, the fatty acid biosynthesis is for the construction of membranes, but without fatty acid degradation, these symbionts are incapable of processing fatty acids as a carbon source. Both N*α*e and N*β*e do not have complete pathways for the creation of glycerophospholipids, yet both contain phospholipid ABC transporters, possibly indicating a dependence on *Nephromyces* for phospholipids.

None of the three endosymbionts contain any of the genes involved in purine degradation (Fig. 2). Both N*β*e and N*α*e do not encode any genes involved in *de novo* purine biosynthesis, including genes for the conversion from IMP to adenine and guanine. Nbe can likely synthesize purines from PPRP though the histidine biosynthesis pathway and contains the genes to synthesize adenine and guanine from IMP. The lack of purine biosynthesis genes in both N*β*e and N*α*e makes these symbionts dependent on *Nephromyces* for both adenine and guanine. If *Nephromyces* were incapable of *de novo* purine biosynthesis, then the entire renal sac community would be dependent on either the tunicate host for all purines or on Nbe. This would likely be a significant burden on the host and adds support to the genomic prediction that *Nephromyces* are able to synthesize purines.

Given their reduced genomes and correspondingly reduced metabolic capabilities, N*β*e and Nbe encode a large proportion of genes for synthesizing amino acids, vitamins, and co-factors (Fig. 2). Together, N*β*e and Nbe could provide *Nephromyces* with all but one essential amino acid (tryptophan). N*α*e is only capable of synthesizing three amino acids, and only one which is an essential amino acid for *Nephromyces* (lysine). Vitamin and cofactor biosynthesis in N *α* eis also limited to synthesizing only heme, ubiquinone, and lipoic acid, with lipoic acid being the only product *Nephromyces* may be incapable of synthesizing itself (Fig. 2). The monophyletic alphaproteobacterium endosymbiont in *Cardiosporidium ciona* also encodes lipoic acid and was predicted to be an essential function (33). None of three bacterial endosymbiont types are functionally equivalent, and FISH microscopy has not revealed any *Nephromyces* cell containing more than a single symbiont type.

No wild *Molgula manhattensis* or *Molgula occidentalis* we examined contained just a single *Nephromyces* species. The average population was three to five *Nephromyces* species in *M. manhattensis* and seven to eight in the larger *M. occidentalis*, with a minimum of two (Fig. 3). From 50 *M. manhattensis* collected from a single locality ~21 species of *Nephromyces* species were recovered. Results were similar for *Molgula occidentalis* 60 individuals, ~19 *Nephromyces* species. Whereas single genus multispecies apicomplexan infections are relatively common (34–37), the extreme diversity and universal multispecies infections found in *Molgula* tunicates is striking. All three bacterial endosymbionts were found in 40% of *M. manhattensis* and *Molgula occidentalis* samples and at least two types were in 90% of *M. manhattensis* and 80% of *Molgula occidentalis*.

Our data shows that *Nephromyces* spp. form complex species swarm communities inside the renal sacs of their hosts, where closely related conspecifics and their functionally distinct bacterial endosymbionts are in immediate proximity with each other. Based on the genomic data from the bacterial endosymbionts, no single bacterial endosymbiont is capable of producing all of the metabolites and precursor molecules that are predicted to be essential for *Nephromyces*. As of yet we have no support of direct metabolite exchange between *Nephromyces* species, but propose that the production of these metabolites may become available in the renal sac indirectly by leaky membranes or cell lysis. This hypothesis does not preclude that these metabolites may also come from the tunicate host, but it is unlikely the expense of an endosymbiont would be tolerated if these metabolites were available in sufficient quantities. The presence of multiple endosymbionts with different types of bacterial endosymbionts may increase the amount and availability of these metabolites inside the renal sac. If this is the case, then multiple *Nephromyces* species may actually decrease *Nephromyces* virulence overall consistent with results from (38). In this way, *Nephromyces* appear to have evolved a complex mutualism with their host, sister species, and bacterial endosymbionts.

## Materials & Methods

### Fluorescence *in situ* Hybridization (F*IS*H) Microscopy

Three 16S targeted bacterial probes with attached fluorophores were used in each fluorescence *in situ* hybridization (F*IS*H) reaction. The probes were designed by Seah 2011 to be specific to the classes of bacterial endosymbionts present in *Nephromyces*. This included alphaproteobacteria (AlexaFluor 555 - 5’ GCATACCGCCAGCGTTCGTT 3’), betaproteobacteria (AlexaFluor 647 - 5’ GCATCCCGCTAGCGTTCAAT 3’), and bacteroides (AlexaFluor 488 - 5’ GCTTGTCGCTAGCGTTTATC 3’) (Seah 2011).

Renal sacks from wild *Molgula manhattensis* collected from Greenwich Bay, Rhode Island, USA, were dissected and emptied onto a positively charged slide. The sample was dried on the slide, and then subjected to a dehydration time series in sterile seawater: 50% ethanol (5 mins), 70% ethanol (8 mins), 95% ethanol (10 mins), and finally 100% ethanol (15 mins). A hybridization solution consisting of 0.9M NaCl, 0.02M Tris-Cl (pH 7.4), 0.01% SDS, 20% HiDi Formamide, and 5% dextran sulfate was mixed with 2pmol/μL of probe, and 200μL applied to the slide (Seah 2011). The slide was then incubated in a humidified dark chamber at 46C overnight. Two washes were conducted by draining the remaining hybridization buffer from the slide and applying 200μL of wash buffer (0.215M NaCl, 0.02M Tris-Cl (pH 7.4), 0.01% SDS) for 30 minutes, with one change of solution (Seah 2011). Samples were then mounted with Prolong Gold w/ DAPI, cured, and imaged using a Zeiss Axioimager M2 imaging system at the URI Genomics and Sequencing Center (URIGSC).

### RNA Extraction & Sequencing

RNA extraction buffer (Zymo Research LLC. Irvine, CA) was added to samples and ground with a pestle. Following grinding, the Zymo Quick-RNA kit (Zymo Research LLC. Irvine, CA) was used and the manufacturer’s protocol was followed. RNA was converted to cDNA and sequenced at the School of Medicine Genome Resource Center, University of Maryland. One paired-end RNA library was run on one lane of the Illumina HiSeq platform. Resulting in 40,606,230 reads from the *M. manhattensis* renal sac.

### RNA Assembly & Binning

Transcriptome data was assembled and proteins were predicted with Trinity/Trinotate pipeline v2.4.0 run on the server at Brown University Center for Computation and Visualization (40). Protein sequences were predicted using Transdecoder (40). BLASTp v 2.8.1 was used to identify bacterial sequences from assembled transcripts against NCBI’s refseq database and binned. Orthofinder was run on remaining Eukaryotic sequences with a custom database of alveolate and ascidian transcriptomes.. Including the following apicomplexan and Chromera transcriptomes downloaded from EuPathDB (*C. parvum* Iowa, *G. niphandrodes, B. bovis* T2Bo, *T. parva* Muguga, *P.falciparum* 3D7, *C. cayetanensis, E. brunetti* Houghton, *E.falciformis* Bayer Haberkorn, *E. tenella* Houghton, *H. hammondi* HH34, *N. caninum* LIV, *S. neurona* SN3, *T. gondii* ME49, *C. velia* CCMP2878, *V.brassicaformis* CCMP3155) and ascidian transcriptomes downloaded form aniseed (*C. intestinalis, H. roretzi, B. schlosseri. M. oculata, M. occulta, M. occidentalis*). Orthologous genes were binned using closest phylogeny in R.

Trimmed reads were mapped back to each of the three bins (*Nephromyces, M. manhattensis, Nephromyces’s* bacteria) and then reassembled independently in Spades (41). *Nephromyces* transcriptome was composed of multiple *Nephromyces* species CDhit v4.6.8 was used to cluster transcripts based on percent identity (42). Transcriptome completeness was assessed with Busco v3 against the Eukaryotic and bacterial reference data sets (43). Transcripts were annotated using Interproscan v71.0 (44).

### Illumina DNA Extraction & Sequencing

The renal sacs from 8 lab grown *M. manhattensis* individuals were dissected and their renal fluid was pooled in a 1.5ml Eppendorf tube. Contents were centrifuged at 8000 g for 5 min. to pellet *Nephromyces* cells, and following centrifugation the renal fluid was discarded. Five hundred microliters of CTAB buffer with 5ul of proteinase K and ceramic beads were added to the pelleted *Nephromyces* cells. The sample was placed in a bead beater for 3 min. and then on a rotator for 1.5hrs at room temp. Five hundred microliters of chloroform were added, mixed gently and centrifuged for 5 min. The top layer was removed and 2x the sample volume of ice-cold 100% EtOH and 10% sample volume of 3M sodium acetate were added to the sample and incubated a −20C overnight. The sample was centrifuged at 16000xg for 30min. and the liquid was removed. Ice cold 70% EtOH was added and centrifuged at 16000xg for 15min. The liquid was removed and the sample air-dried for 2 min. DNA was re-eluted in 50ul of deionized water. A nanodrop (2000c, ThermoScientific) was used to assess DNA purity and DNA concentration, and a genomic gel was run to assess DNA fragmentation. Following quality control, an Illumina library was constructed. Library prep and sequencing were done at the URI Genomics and Sequencing Center (URIGSC). The completed library was sequenced on the Illumina Miseq platform at the URIGSC and the HiSeq platform at the University of Maryland Baltimore sequencing center on three lanes.

### Illumina Assembly

One MiSeq lane and three lanes of HiSeq, all from the same library, were trimmed using Trimmomatic v0.39 (45) then assembled using SPAdes v3.13.0 (41) assembler on the URI server BlueWaves.

### Pacific Biosciences DNA Extraction & Sequencing

Using the contents of 150 (done in batches of 10 then pooled) *M. manhattensis* renal sacs, the same DNA extraction protocol was performed as for Illumina sequencing. DNA was sequenced using three SMRT cells on the Pacific Biosciences platform at the University of Baltimore sequencing center.

### Pacific Biosciences Assembly

Pacific Biosciences reads were error corrected using PBSuite v15.8.24 (46) on the Brown University server, Oscar. Reads were then assembled using Canu v1.6 (47). Contigs generated by Canu were combined with Illumina MiSeq/HiSeq short reads with ABySS v2.02 (48). *Nephromyces* contigs were identified by mapping *Nephromyces* transcriptome reads to the genomic assembly using Bowtie2 v1.2.2. Contigs with greater than 90x coverage assessed by bedtools (49) were binned as *Nephromyces* additional screening of binned contigs was done using VizBin (50). Contigs were additionally binned using CAT (51), and those that were classified as apicomplexan were added to the existing assembly.

### Bacterial Endosymbiont Genome Assembly

Using the contigs from the ABySS assembly bacterial contigs were initially identified by hexamers using VizBin, transcriptomic reads that were identified as bacterial were mapped using Bowtie2 (52). Bacterial contigs were separated based on a 90x coverage threshold with bedtools (49). Binned bacterial contigs were preliminarily annotated with Prokka (53). Resulting annotations were run through KEGG GhostKoala to assign and separate by taxonomy. Taxon separated contig bins were merged and scaffolded using PBJelly from the PBsuite v15.8.24 of tools (54). Illumina MiSeq and HiSeq reads were remapped to resulting contigs to ensure accurate assembly using Bowtie2. Final assembled bacterial genomes were re-annotated with Prokka using a genus-specific database.

### Data

The pangenome and transcriptome of *Nephromyces*, as well the associated bacterial endosymbionts, are available on GenBank under the bioproject ID PRJNA666913.

### Abridged Amplicon Methods

Fifty *Molgula manhattensis* tunicates were collected from a single floating dock located in Greenwich Bay, RI (41.653N, −71.452W), and 54 *Molgula occidentalis* were collected from Alligator Harbor, FL (29.899N, −84.381W) by Gulf Specimens Marine Laboratories, Inc. (https://gulfspecimen.org/). DNA was extracted with previously described methods. *Nephromyces* specific COI primers and universal 16S rRNA primers (55) were PCR amplified, and sequenced on two Illumina Miseq runs (2×250 2×200). Resulting sequencing reads were error corrected and merged using dada2 with pool=sudo (56). Amplicon sequence variants (ASVs) were stringently filtered to only ASVs with greater than 20 reads per sample. *Nephromyces* COI ASVs were clustered based on percent identity from 100-94% with CD-hit. For full methods see supplemental materials.

### *Molgula manhattensis* laboratory culture

*Molgula manhattensis* tunicates were collected from a dock in Greenwich Bay, Rhode Island (41.653N, −71.452W) during the summer of 2014. In batches of five Gonads were dissected from sexually mature, *M. manhattensis*. Eggs and sperm were mixed with sterile seawater and divided evenly between two petri dishes. This was repeated 4 times for a total of eight plates. Plates were incubated at room temperature for two days with daily 100% water changes. Tunicate larvae attached to the bottom and sides of the petri dishes by day three. By day four, larvae had metamorphosed into adults and were actively feeding. Plates were moved to an incubator at 18° C with a 24 hr dark cycle to limit growth of contaminants. Tunicates were fed by 100% water exchange with cultures of *Isochrysis galbana* and *Chaetoceros gracilis* three days a week. After ten days a single *M. manhattensis* was dissected and the renal fluid was extracted with a syringe and placed in a 1.5ml tube. 1μ or renal fluid was added to two plates, 1μ of renal fluid diluted 1/100 was added to two plates, and a single oocyst was picked and added to two plates, and two plates were kept *Nephromyces* free. After several weeks tunicates were moved to aerated beakers to meet their increased nutrient and gas exchange requirements. Feeding regimen remained the same except that food volume was increased with tunicate growth. Tunicates were grown for six months until they were ~10mm across. After 6 months tunicates were periodically sacrificed to cheek infection status

## References

1. C. M. Cavanaugh, S. L. Gardiner, M. L. Jones, H. W. Jannasch, J. B. Waterbury, Prokaryotic Cells in the Hydrothermal Vent Tube Worm Riftia pachyptila Jones: Possible Chemoautotrophic Symbionts. Science 213, 340–342 (1981).

2. A. Orsi, et al., Dynamic and transient interactions of Atg9 with autophagosomes, but not membrane integration, are required for autophagy. Mol. Biol. Cell 23, 1860–1873 (2012).

3. M. B. Saffo, Nitrogen Waste or Nitrogen Source? Urate Degradation in the Renal Sac of Molgulid Tunicates. Biol. Bull. 175, 403–409 (1988).

4. M. B. Saffo, Symbiosis within a symbiosis: Intracellular bacteria within the endosymbiotic protistNephromyces. Mar. Biol. 107, 291–296 (1990).

5. M. B. Saffo, A. M. McCoy, C. Rieken, C. H. Slamovits, Nephromyces, a beneficial apicomplexan symbiont in marine animals. Proc. Natl. Acad. Sci. 107, 16190–16195 (2010).

6. C. Paight, C. H. Slamovits, M. B. Saffo, C. E. Lane, *Nephromyces* Encodes a Urate Metabolism Pathway and Predicted Peroxisomes, Demonstrating That These Are Not Ancient Losses of Apicomplexans. Genome Biol. Evol. 11, 41–53 (2019).

7. A. A. Escalante, F. J. Ayala, Evolutionary origin of Plasmodium and other Apicomplexa based on rRNA genes. Proc. Natl. Acad. Sci. 92, 5793–5797 (1995).

8. N. A. Counihan, M. Kalanon, R. L. Coppel, T. F. de Koning-Ward, Plasmodium rhoptry proteins: why order is important. Trends Parasitol. 29, 228–236 (2013).

9. R. Cardoso, H. Soares, A. Hemphill, A. Leitão, Apicomplexans pulling the strings: manipulation of the host cell cytoskeleton dynamics. Parasitology 143, 957–970 (2016).

10. A. J. Reid, Large, rapidly evolving gene families are at the forefront of host-parasite interactions in *Apicomplexa*. Parasitology 142, S57–S70 (2015).

11. J. Janouškovec, et al., Factors mediating plastid dependency and the origins of parasitism in apicomplexans and their close relatives. Proc. Natl. Acad. Sci. 112, 10200–10207 (2015).

12. Y. H. Woo, et al., Chromerid genomes reveal the evolutionary path from photosynthetic algae to obligate intracellular parasites. eLife 4 (2015).

13. M. Zarowiecki, M. Berriman, What helminth genomes have taught us about parasite evolution. Parasitology 142, S85–S97 (2015).

14. S. A. Frank, Host-Symbiont Conflict over the Mixing of Symbiotic Lineages. Proc. Biol. Sci. 263, 339–344 (1996).

15. S. Alizon, A. Hurford, N. Mideo, M. Van Baalen, Virulence evolution and the trade-off hypothesis: history, current state of affairs and the future: Virulence evolution and trade-off hypothesis. J. Evol. Biol. 22, 245–259 (2009).

16. C. E. Cressler, D. V. McLEOD, C. Rozins, J. Van Den Hoogen, T. Day, The adaptive evolution of virulence: a review of theoretical predictions and empirical tests. Parasitology 143, 915–930 (2016).

17. S. A. Frank, Models of Parasite Virulence. Q. Rev. Biol. 71, 37–78 (1996).

18. M. B. Saffo, R. Nelson, The cells of *Nephromyces*: developmental stages of a single life cycle. Can. J. Bot. 61, 3230–3239 (1983).

19. S. A. Muñoz-Gómez, et al., Nephromyces Represents a Diverse and Novel Lineage of the Apicomplexa That Has Retained Apicoplasts. Genome Biol. Evol. 11, 2727–2740 (2019).

20. J. R. Nolfi, Biosynthesis of uric acid in the tunicate, Molgula manhattensis, with a general scheme for the function of stored purines in animals. Comp. Biochem. Physiol. 35, 827–842 (1970).

21. C. C. Lambert, G. Lambert, G. Crundwell, K. Kantardjieff, Uric Acid Accumulation in the Solitary Ascidian. 10 (1998).

22. M. A. Rouf, R. F. Lomprey, Degradation of Uric Acid by Certain Aerobic Bacteria. J. Bacteriol. 96, 617–622 (1968).

23. A. Thong-On, et al., Isolation and Characterization of Anaerobic Bacteria for Symbiotic Recycling of Uric Acid Nitrogen in the Gut of Various Termites. Microbes Environ. 27, 186–192 (2012).

24. W. J. Middelhoven, G. S. De Hoog, S. Notermans, Carbon assimilation and extracellular antigens of some yeast-like fungi. Antonie Van Leeuwenhoek 55, 165–175 (1989).

25. J. F. Yamagishi, N. Saito, K. Kaneko, The advantage of leakage of essential metabolites and resultant symbiosis of diverse species. Phys. Rev. Lett. 124, 048101 (2020).

26. N. A. Moran, H. E. Dunbar, J. L. Wilcox, Regulation of Transcription in a Reduced Bacterial Genome: Nutrient-Provisioning Genes of the Obligate Symbiont Buchnera aphidicola. J. Bacteriol. 187, 4229–4237 (2005).

27. M. J. López-Sánchez, et al., Evolutionary Convergence and Nitrogen Metabolism in Blattabacterium strain Bge, Primary Endosymbiont of the Cockroach Blattella germanica. PLoS Genet. 5, e1000721 (2009).

28. D. J. de Souza, A. Bézier, D. Depoix, J.-M. Drezen, A. Lenoir, Blochmannia endosymbionts improve colony growth and immune defence in the ant Camponotus fellah. BMC Microbiol. 9, 29 (2009).

29. H. Urakawa, et al., Hydrothermal vent gastropods from the same family (Provannidae) harbour e- and gamma-proteobacterial endosymbionts. Environ. Microbiol. 7, 750–754 (2005).

30. B. Marin, E. C. M. Nowack, M. Melkonian, A Plastid in the Making: Evidence for a Second Primary Endosymbiosis. Protist 156, 425–432 (2005).

31. B. Seah, “A Tripartite Animal-Protist-Bacteria Symbiosis: Culture-Independent and Phylogenetic Characterization.” (2011).

32. Y. Nishimoto, et al., Evolution and phylogeny of the heterogeneous cytosolic SSU rRNA genes in the genus Plasmodium. Mol. Phylogenet. Evol. 47, 45–53 (2008).

33. Elizabeth S. Hunter, Christopher Paight, Christopher E. Lane, Metabolic contributions of an alphaproteobacterial endosymbiont in the apicomplexan Cardiosporidium cionae. Front. Microbiol. (2020).

34. T. J. C. Anderson, et al., Microsatellite Markers Reveal a Spectrum of Population Structures in the Malaria Parasite Plasmodium falciparum. Mol. Biol. Evol. 17, 1467–1482 (2000).

35. K.-S. Lee, et al., Plasmodium knowlesi: Reservoir Hosts and Tracking the Emergence in Humans and Macaques. PLoS Pathog. 7, e1002015 (2011).

36. A. Arnott, A. E. Barry, J. C. Reeder, Understanding the population genetics of Plasmodium vivax is essential for malaria control and elimination. Malar.J. 11, 14 (2012).

37. A. Lalremruata, et al. Species and genotype diversity of Plasmodium in malaria patients from Gabon analysed by next generation sequencing. Malar.J. 16 (2017).

38. P. Nelson, G. May, Defensive Symbiosis and the Evolution of Virulence. Am. Nat. 196, 333–343 (2020).

39. Y.-L. Lau, et al., Deciphering the Draft Genome of Toxoplasma gondii RH Strain. PLoS ONE 11 (2016).

40. B. J. Haas, et al., De novo transcript sequence reconstruction from RNA-seq using the Trinity platform for reference generation and analysis. Nat. Protoc. 8, 1494–1512 (2013).

41. A. Bankevich, et al., SPAdes: A New Genome Assembly Algorithm and Its Applications to Single-Cell Sequencing. J. Comput. Biol. 19, 455–477 (2012).

42. W. Li, A. Godzik, Cd-hit: a fast program for clustering and comparing large sets of protein or nucleotide sequences. Bioinformatics 22, 1658–1659 (2006).

43. F. A. Simão, R. M. Waterhouse, P. Ioannidis, E. V. Kriventseva, E. M. Zdobnov, BUSCO: assessing genome assembly and annotation completeness with single-copy orthologs. Bioinformatics 31, 3210–3212 (2015).

44. R. D. Finn, et al., InterPro in 2017—beyond protein family and domain annotations. Nucleic Acids Res. 45, D190–D199 (2017).

45. A. M. Bolger, M. Lohse, B. Usadel, Trimmomatic: a flexible trimmer for Illumina sequence data. Bioinformatics 30, 2114–2120 (2014).

46. A. C. English, et al., Mind the Gap: Upgrading Genomes with Pacific Biosciences RS Long-Read Sequencing Technology. PLoS ONE 7, e47768 (2012).

47. S. Koren, et al., Canu: scalable and accurate long-read assembly via adaptive *k*-mer weighting and repeat separation. Genome Res. 27, 722–736 (2017).

48. S. D. Jackman, et al., ABySS 2.0: resource-efficient assembly of large genomes using a Bloom filter. Genome Res. 27, 768–777 (2017).

49. A. R. Quinlan, I. M. Hall, BEDTools: a flexible suite of utilities for comparing genomic features. Bioinformatics 26, 841–842 (2010).

50. C. C. Laczny, et al., VizBin - an application for reference-independent visualization and human-augmented binning of metagenomic data. Microbiome 3, 1 (2015).

51. F. A. B. von Meijenfeldt, K. Arkhipova, D. D. Cambuy, F. H. Coutinho, B. E. Dutilh, Robust taxonomic classification of uncharted microbial sequences and bins with CAT and BAT. bioRxiv (2019) https:/doi.org/10.1101/530188 (April 4, 2019).

52. B. Langmead, S. L. Salzberg, Fast gapped-read alignment with Bowtie 2. Nat. Methods 9, 357–359 (2012).

53. T. Seemann, Prokka: rapid prokaryotic genome annotation. Bioinformatics 30, 2068–2069 (2014).

54. B. Bushell, BBMap. SourceForge (2014) (August 11, 2020).

55. A. Klindworth, et al. Evaluation of general 16S ribosomal RNA gene PCR primers for classical and next-generation sequencing-based diversity studies. Nucleic Acids Res. 41, e1–e1 (2013).

56. B. J. Callahan, et al., DADA2: High resolution sample inference from Illumina amplicon data. Nat. Methods 13, 581–583 (2016).

57. M. D. Lee, GToTree: a user-friendly workflow for phylogenomics. Bioinformatics 35, 4162–4164 (2019).

58. S. R. Eddy, Accelerated Profile HMM Searches. PLoS Comput. Biol. 7, e1002195 (2011).

59. S. Capella-Gutiérrez, J. M. Silla-Martínez, T. Gabaldón, trimAl: a tool for automated alignment trimming in large-scale phylogenetic analyses. Bioinformatics 25, 1972–1973 (2009).

60. D. Hyatt, et al., Prodigal: prokaryotic gene recognition and translation initiation site identification. BMC Bioinformatics 11, 119 (2010).

61. R. C. Edgar, MUSCLE: a multiple sequence alignment method with reduced time and space complexity. BMC Bioinformatics 5, 113 (2004).

62. W. Shen, J. Xiong, “TaxonKit: a cross-platform and efficient NCBI taxonomy toolkit” (Bioinformatics, 2019) https:/doi.org/10.1101/513523 (October 7, 2020).

63. M. N. Price, P. S. Dehal, A. P. Arkin, FastTree 2 – Approximately Maximum-Likelihood Trees for Large Alignments. PLoS ONE 5, e9490 (2010).

64. G. M. Boratyn, J. Thierry-Mieg, D. Thierry-Mieg, B. Busby, T. L. Madden, Magic-BLAST, an accurate DNA and RNA-seq aligner for long and short reads. bioRxiv (2018) https:/doi.org/10.1101/390013 (February 24, 2019).

65. M. Martin, Cutadapt removes adapter sequences from high-throughput sequencing reads. EMBnet.journal 17, 10–12 (2011).

66. L. Guillou, et al, The Protist Ribosomal Reference database (PR2): a catalog of unicellular eukaryote Small Sub-Unit rRNA sequences with curated taxonomy. Nucleic Acids Res. 41, D597–D604 (2012).

67. K. Katoh, D. M. Standley, MAFFT Multiple Sequence Alignment Software Version 7: Improvements in Performance and Usability. Mol. Biol. Evol. 30, 772–780 (2013).

68. H. Wickham, Create Elegant Data Visualisations Using the Grammar of Graphics (2016) (September 3, 2020).

69. S. J. Booker, Unraveling the Pathway of Lipoic Acid Biosynthesis. Chem. Biol. 11, 10–12 (2004).

