## Supplemental Material for "Codependence in the *Nephromyces* species swarm depends on heterospecific bacterial endosymbionts"

### Supplementary Materials - Methods

Bacterial Phylogeny

Using the predicted phylogenetic positions from the Microbial Gene Atlas (MiGA), all complete genomes for the classes betaproteobacteria, alphaproteobacteria, and Bacteroides available on NCBI were collected. The GToTree pipeline was run on each of these datasets, including the *Nephromyces* endosymbiont, using the relevant HMM set of single copy gene targets (57–62). This included 138 gene targets and 722 genomes in alphaproteobacteria, 203 gene targets and 471 genomes in betaproteobacteria, and 90 gene targets and 388 genomes in Bacteroidetes. In betaproteobacteria, 5 genomes were removed for having either too few hits to the single copy gene targets, or multiple hits. The final trees were created with FastTree v2 (63), and formatted in FigTree (S Figure 2,3).

Amplicon Methods Detailed

Fifty *Molgula manhattensis* tunicates were collected from a single floating dock located in Greenwich Bay, RI (41.653N, -71.452W), and 29 *Molgula occidentalis* were collected from Alligator Harbor, FL (29.899N, -84.381W) by Gulf Specimens Marine Laboratories, Inc. (https://gulfspecimen.org/). All 79 samples were collected in August of 2016 and prepared for a single Illumina MiSeq flow cell (hereafter referred to as Run One). An additional 25 *Molgula occidentalis* were collected by Gulf Specimens Marine Laboratories, Inc. in March of 2018 from the same location and prepared for a second MiSeq flow cell (hereafter referred to as Run Two). Tunicates were dissected to remove renal sacs and *Nephromyces* cells contained within were collected by a micropipette and placed in 1.5 ml eppendorf tubes. Dissecting tools were sterilized in a 10% bleach solution for 15 min and then rinsed between tunicates. Sample tubes were immediately frozen in liquid nitrogen for five minutes and subsequently stored at -80°C.

DNA was extracted using the method described in (6) and stored at -20° C. The 18S rRNA primers (Run One only) and CO1 primers were designed to target *Nephromyces* based on available genomic data. The universal 16S rRNA primers from (55) were used to amplify the bacterial endosymbionts within *Nephromyces*. The Illumina adaptor sequence was added to the start of each primer resulting in the following sequences 18Sf (TCGTCGGCAGCGTCAGATGTGTATAAGAGACAGCGGTAATTCCAGCTCC), 18Sr (GTCTCGTGGGCTCGGAGATGTGTATAAGAGACAGTGCTTTCGCAGTAGTYYGTCTTT), CO1f (TCGTCGGCAGCGTCAGATGTGTATAAGAGACAGYGGWGTAGGWSCWGGWTGGA), CO1r (GTCTCGTGGGCTCGGAGATGTGTATAAGAGACAGACTTCWGGATGWCCAAARAA) 16Sf (TCGTCGGCAGCGTCAGATGTGTATAAGAGACAGCCTACGGGNGGCWGCAG), 16Sr (GTCTCGTGGGCTCGGAGATGTGTATAAGAGACAGGACTACHVGGGTATCTAATCC). PCR was performed for each sample for all three primer sets with the following cycle 94° C 2 min (94° C 30 sec, 55° C 30 sec, 72° C 45 sec) x 35, 72° C 5 min. PCR product was visually inspected on an agarose gel and quantity estimated with a nanodrop. For Run One, 20 ml of PCR product from each of the three primer sets were pooled into a single tube corresponding to each individual tunicate. For Run Two samples, 16S and COI products were sequenced separately because of the lower number of samples compared to Run One. The PCR products for both runs were cleaned using ampure bead purification (Beckman Coulter) with a 0.7 percent solution. The addition of well specific adaptors, library preparation, and sequencing was done at the URI genome sequencing center on the Illumina MiSeq platform using the Illumina MiSeq reagent kit V2 2X250.

Run One sequences were de-multiplexed prior to analysis and BBDuck, from the BBMap suite of tools, was used to bin reads based on CO1 primers (54). The universal 18S rRNA and 16S rRNA primers were too conserved for reliable binning based on primers, so reads were screened against the PR2 database using the NCBI's magicblast (64). Sequences with a 85% identity and 35% coverage were classified as 18S sequences and binned into a new file composed of 18S reads. Adaptors and primer sequences were removed from the forward and reverse reads from each of the three read sets using BBDuck. Since Run Two was not multiplexed, adaptors and primers were removed with cutadapt v2.10 (65).

Cleaned and binned read sets were individually processed in R using dada2 with the pool="pseudo" setting (56). Assembled 18S and 16S were assigned taxonomies with the PR2 database (66). CO1 sequences were assigned taxonomy using blastx against NCBI's refseq protein database. All 18S and CO1 sequences that did not return an apicomplexan as top hit were removed from the count table, taxonomy table, and sequence files. The 16S sequences were aligned with MAFFT (67) to 16S rRNA sequences from the three known bacterial endosymbionts found in *Nephromyces,* an alphaproteobacteria, betaproteobacteria and Bacteroides. Reference sequences were trimmed to the amplicon sequence length and CDhit was used to cluster sequences with 85% sequence identity (42). All bacterial sequences, which did not cluster were deemed contamination and removed from the count table, taxonomy table, and sequences file. Samples with less than 5% of the mean number of reads were considered failed and were removed (three samples). Any ASV represented by less than 20 reads from a single sample were set to 0.

The remaining 18S and CO1 sequences were aligned with MAFFT and trimmed to the same length. Sequences were clustered at 100%, 99%, 98%, 97%, 96%, 95%, 94% sequence identity levels using CDhit. The 18S, CO1 clusters and 16S bins corresponding to endosymbiont type were processed in R. Figures were made in R using ggplot (68).

In order to estimate clustering thresholds for species assignment from ASVs, all available *Plasmodium* COI sequences were downloaded from NCBI's Genbank. Sequences were aligned with MAFFT and trimmed to the region amplified by our COI primers. Sequences were then clustered with CD-hit at 99, 98, 97, 96% sequence identity. Clustered at 98% sequence identity collapsed almost all of the sequences to assigned species levels without collapsing multiple species. At 97% all sequences collapse to species level and a few species are collapsed. We clustered *Nephromyces* along the same gradient (Fig 3), and based on the *Plasmodium* analysis, the 98% and 97% sequence identity ASV cluster for *Nephromyces*.

### Supplementary Materials - Results

#### *Nephromyces* Bacterial Endosymbiont Genomic Characterization

##### *Nephromyces's* Alphaproteobacterial genome (N $\alpha$ e)

Multiple alphaproteobacterial endosymbionts were recovered from our genomic data and assembled into a draft genome. The presence of multiple closely related alpha proteobacteria genomes with high AT bias (25% GC content) and regions of low complexity have limited our ability to assemble these genomes completely. The two genomes assemble into 11 contigs ranging in size from 13 kb to 312 kb for a combined length of 995,540 and an average of 90,503 (S Figure 4). The Microbial Gene Atlas (MiGA) predicted this assembly is 85.6% complete with 0% contamination. The draft genome contains 844 predicted coding sequences, 35 tRNAs matching all codons, and 4 rRNAs. 546 of the predicted genes have KASS annotations. (119 Genetic information processing, 32 Carbohydrate metabolism, 30 Energy metabolism, 29 Cellular processes, 26 Nucleotide metabolism, 26 Metabolism of cofactors and vitamins, 25 environmental information processing, 15 lipid metabolism, 12 amino acid metabolism, 15 unclassified).

Nephromyces's Bacteroidetes Genome (NBe)

*Nephromyces* bacteroidetes genome is circular, 494,352 nucleotides long and extremely AT rich (22% GC content) (S Figure 4). The genome contains 503 predicted genes, 31 tRNAs predicted to recognize all codons, and 4 rRNAs. 391 of the predicted genes have KASS annotations. (110 Genetic information processing, 40 Carbohydrate metabolism, 38 Energy metabolism, 31 amino acid metabolism, 21 Metabolism of cofactors and vitamins, 17 Nulcleotide metabolism, 11 unclassified, 10 Lipid metabolism, 9 Cellular processes) Nephromyces's Betaproteobacterial Genome (N $\beta$ e) The Betaproteobacterial genome is circular 866,396 bp long with 30%GC content. It contains 880 predicted genes, 40 tRNAs, and 4 rRNAs (two identical 16s copies and 2 identical 23s copies). 753 of the 880 predicted genes have KAAS annotations (156 Genetic information processing, 61 Carbohydrate metabolism, 47 Energy metabolism, 11 Cellular processes, 45 Nucleotide metabolism, 68 Metabolism of cofactors and vitamins, 39 environmental information processing, 18 lipid metabolism, 62 amino acid metabolism, 14 unclassified).

*Nephromyces's* Betaproteobacterial Genome (N $\beta$ e)

The Betaproteobacterial genome is circular 866,396 bp long with 30% GC content (S Figure 4). It contains 880 predicted genes, 40 tRNAs, and 4 rRNAs (two identical 16s copies and 2 identical 23s copies). 753 of the 880 predicted genes have KAAS annotations (156 Genetic information processing, 61 Carbohydrate metabolism,47 Energy metabolism, 11 Cellular processes, 45 Nucleotide metabolism, 68 Metabolism of cofactors and vitamins, 39 environmental information processing,18 lipid metabolism, 62 amino acid metabolism, 14 unclassified).

Bacterial Phylogeny

The *Nephromyces* alphaproteobacteria was shown to be sister to the genus *Rickettsia* (33). The *Nephromyces* betaproteobacteria endosymbiont was shown to fall within the Alcaligenaceae family, sister to another group of endosymbionts *Candidatus* Kinetoplastibacterium, endosymbionts of trypanosomes. Similarly, the bacteroides endosymbiont fell within the Flavobacteriaceae family, sister to *Candidatus* Sulcia muelleri, *Blattabacterium, Candidatus* Uzinura diaspidicola, and *Candidatus* Walczuchella monophlebidarum. All of these species are bacterial endosymbiont of insects. This is likely a result of the characteristic convergent genome reduction of vertically transmitted bacterial endosymbionts.

### *Nephromyces* Metabolic Pathway Characterization

#### Endocytosis

*Nephromyces* encodes genes for clathrin-dependant endocytosis. An additional 25 genes related to endocytosis are predicted in *Nephromyces* than in *Plasmodium falciparum.* Many of the additional genes found in *Nephromyces* are in the VPS and CHMP protein families, which form part of the ESCRT machinery and are important in the biogenesis of multivesicular bodies. Multivesicular bodies transport ubiquitinated proteins to lysosomes for degradation. Other proteins not found in *P. falciparum* include a number of genes in the AP2 complex, which are accessory proteins in clathrin-mediated endocytosis. The AP2 complex plays an important role in the regulation of the assembly of clathrin-coated vesicles.

#### Carbohydrate Metabolism

Basic carbon metabolism in *Nephromyces* is similar to other apicomplexans. It encodes the complete pathways for the citric acid cycle, glycolysis, gluconeogenesis, and pentose phosphate pathway. Interestingly, *Nephromyces* encodes far more genes involved with inositol phosphate metabolism than either *P. falciparum* or *Toxoplasma gondii.* Despite the absence of genes involved in the synthesis of myo-inositol in *P. falciparum* or *T. gondii*, there is support that myo-inositol is used in intracellular calcium signaling in these two organisms. How *P. falciparum* or *T. gondii* is able to use myo-inositol without being able to synthesize it is unclear; it is presumed that they have divergent and unrecognizable myo-inositol biosynthesis genes. However, *Nephromyces* is predicted to be able to synthesize myo-inositol, and these genes are readily identifiable as orthologous to myo-inositol biosynthesis genes in other organisms. *Nephromyces* also has a copy of serine-pyruvate aminotransferase (AGXT), which catalyzes the conversion of glyoxylate to glycine and pyruvate. Additionally, *Nephromyces* encodes malate synthase, which in combination with acetyl-CoA forms malate from glyoxylate (6).

#### Fatty Acid Metabolism

*Nephromyces* is able to perform fatty acid initiation and elongation in both the mitochondrial and cytoplasmic pathways, as well as elongation in the endoplasmic reticulum. Additionally, *Nephromyces* encodes D-glycerate 3-kinase (GLYK), aldehyde dehydrogenase (ALDH), alcohol dehydrogenase (ADH), glycerol kinase (glpK), glycerol-3-phosphate O-acyltransferase (GPAT1), and 1-acyl-sn-glycerol-3-phosphate acyltransferase (plsC) and is thus able to create triglycerides from glucose.

Nucleic acid metabolism

*Nephromyces* is unique among apicomplexans because it contains a complete pathway for the biosynthesis of inosine monophosphate (IMP). IMP is a purine and the starting molecule for the biogenesis of guanine and adenine. *De novo* biosynthesis of purines has been lost in all sequenced apicomplexans. Other apicomplexans are capable of scavenging precursor molecules to IMP and converting them into IMP, but *Nephromyces/Cardiosporidium* encodes the entire IMP biosynthesis pathway from 5-Phosphoribosyl diphosphate (PPRP). The genes involved in this pathway include amidophosphoribosyltransferase (purF), phosphoribosylamine---glycine ligase (purD), phosphoribosylformylglycinamidine synthase (purL), phosphoribosylformylglycinamidine cyclo-ligase (purM), phosphoribosylaminoimidazole carboxylase (PAICS), phosphoribosylaminoimidazole-succinocarboxamide synthase (purC), adenylosuccinate lyase (purB), IMP cyclohydrolase (purH). *Nephromyces* encodes for the metabolic machinery for the biosynthesis of adenine and guanine from the end product of this pathway, IMP. In addition to the biosynthesis of purines, *Nephromyces* are also the only known apicomplexans capable of purine degradation (6).

Biosynthesis of Amino Acids

*Nephromyces* is predicted to be able to synthesize 10 amino acids including alanine, asparagine, aspartic acid, cysteine, glutamine, glutamic acid, glycine, methionine, serine, and threonine. In addition to the complete pathways, *Nephromyces* has partial pathways for the synthesis of phenylalanine from phenylpyruvate and can convert tyrosine from phenylalanine. *Nephromyces* also encodes branched-chain amino acid aminotransferase, which adds the final amine group to valine, leucine, and isoleucine.

Vitamin and cofactor synthesis

*Nephromyces* has the predicted biosynthetic capabilities to produce riboflavin, acetyl CoA, nicotinate, folate, retinol, vitamin E, heme, and ubiquinone. *Nephromyces* encodes a copy of lipoyl synthase (lipA), but lacks lipoyl(octanoyl) transferase (lipB) in the lipoic acid synthesis pathway. Lipoic acid is an essential cofactor involved in the citric acid cycle, pyruvate dehydrogenase complex, 2-oxoglutarate dehydrogenase complex, branched-chain oxoacid dehydrogenase, and acetoin dehydrogenase (69).

Endosymbiont Metabolic Pathway Characterization

Carbohydrate metabolism

The $\alpha$-proteobacterial endosymbiont of *Nephromyces* (N$\alpha$e) has an extremely reduced carbohydrate metabolism. Including all of the genes involved with gluconeogenesis and glycolysis. The only carbohydrate metabolism genes present are a complete citrate acid cycle and pyruvate dehydrogenase E1 component (aceE), dihydrolipoamide dehydrogenase (pdhD), and pyruvate dehydrogenase E2 component (aceF), which converts pyruvate to acetyl-CoA. With such severe reduction in carbohydrate metabolism, pyruvate appears to be the only carbon source the alphaproteobacteria is capable of processing.

The bacteroidetes endosymbiont (Nbe) has a similarly reduced carbohydrate metabolism as in the $\alpha$-proteobacteria, however the reduction is not as extreme. Having lost gluconeogenesis the bacteroidetes endosymbiont can process fructose into phosphoenolpyruvate and contains pyruvate dehydrogenase E1 component (aceE), dihydrolipoamide dehydrogenase (pdhD), and pyruvate dehydrogenase E2 component (aceF) to convert pyruvate into acetyl-CoA. While the full citrate cycle is incomplete the partial cycle from 2-oxoglutarate to oxaloacetate is complete, as well as the reductive pentose phosphate pathway from glyceraldehyde-3P to ribulose-5P.

*Nephromyces's* b-proteobacterial endosymbiont (Ne) has the most complete carbohydrate metabolism encoding the complete non-oxidative pentose phosphate pathway, the citrate cycle from 2-oxoglutarate to oxaloacetate, and the core glycolysis module involving three carbon compounds.

Fatty Acid metabolism

Paradoxically, while the enzymes involved with fatty acid biosynthesis (from malonyl-CoA in N$\alpha$e and N$\beta$e and from acetyl-CoA in Nbe) are present in all three types of bacterial endosymbiont, all the genes involved in fatty acid degradation have been lost in every symbiont. N$\alpha$e and N$\beta$e have a reduced glycerophospholipid metabolism and must convert phosphatidate to synthesis phosphatidylethanolamine, phosphatidylglycerol, and phosphatidylserine, while N$\beta$e is able to *de novo* synthesize glycerophospholipids from glycerone.

Nucleic acid metabolism

Purine metabolism is similarly reduced in both N$\alpha$e and Nbe. Both endosymbionts lack all genes in the IMP biosynthesis pathway, as well as the ability to convert IMP to guanine or adenine. With so few purine biosynthesis capabilities, all of the guanine and adenine for DNA replication must be obtained as preformed nucleobases. N$\beta$e purine biosynthesis is complete encoding adenylosuccinate lyase (purB), and IMP cyclohydrolase (purH) and is able to synthesize IMP from 5-Phosphoribosyl diphosphate (PRPP) (with the histidine synthesis pathway) as well as the genes required to convert IMP to both guanine and adenine. However, none of the three endosymbionts encode any genes involved in purine degradation.

Both Ne and N$\beta$e encode the necessary genes for pyrimidine biosynthesis. However, because Nbe is missing several genes involved in pyrimidine biosynthesis it seems likely that Nbe is dependent on *Nephromyces* for both purines and pyrimidines

Biosynthesis of Amino Acids

N$\alpha$e is only capable of synthesizing the amino acids glutamine, glutamic acid, and lysine. Of these three amino acids, lysine is the only amino acid that *Nephromyces* is unable to synthesize.

Nbe is capable of synthesizing 11 amino acids: arginine, aspartic acid, glutamine, glutamic acid, glycine, isoleucine, leucine, lysine, serine, threonine, and valine. Bacteroidetes is also able to synthesis phenylpyruvate, but lack the ability to synthesis phenylalanine. Biosynthesis of arginine, isoleucine, leucine, lysine, and valine are not present in the *Nephromyces* transcriptome and may represent the bacteroidetes contribution. Additionally, Bacteroidetes synthesis of phenylpyruvate, but inability to synthesis phenylalanine compliments *Nephromyces* synthesis of phenylalanine from phenylpyruvate, but inability to synthesis phenylpyruvate.

N $\beta$ e encodes the genes for 11 amino acids: arginine, glutamine, glutamic acid, glycine, histidine, lysine, phenylalanine, proline, serine, threonine, and tyrosine. N $\beta$ e also encodes all the genes for synthesis of isoleucine, leucine, and valine, but lacks the last gene in the pathway branched-chain amino acid aminotransferase. However, *Nephromyces* encodes branched-chain amino acid aminotransferase and may be able to complete isoleucine, leucine, and valine by adding the final amine group.

Vitamin and cofactor synthesis

N $\alpha$ e only encodes genes for the biosynthesis three vitamins and co-factors; heme, ubiquinone, and Lipoic acid. As lipoic acid can only be partially synthesized in *Nephromyces.* Lipoic acid may be an important product produced by the alphaproteobacteria. In contrast Nbe is capable of biosynthesis of vitamin B6, lipoic acid, folate and vitamin K. N $\beta$ e is able to synthesize most cofactors including riboflavin, vitamin B6, nicotinate, coenzyme-A, folate, heme, and ubiquinone.

Secretion and Transporters

All three symbionts encode at least a partial bacterial sec secretion system. N $\beta$ e is the most complete missing only secM. N $\alpha$ e lacks secE, secG, and secM. Nbe is the most incomplete missing secB, secD, secE, secM with so many genes missing it is not clear if the sec secretion system is functional in Nbe. Both N $\beta$ e and Nbe have the twin-arginine translocation pathway. N $\alpha$ e has TatC, but lacks TatA and is therefore incomplete. While N $\alpha$ e is missing the Tat transport system, N $\alpha$ e does encode the type 4 bacterial secretion system.

In addition to the more general secretion systems some more specific ABC transporters. Nbe has the fewest ABC transporters only phospholipid and possibly a heme transporter. N $\alpha$ e also encodes a phospholipid and heme transporter in addition to a lipoprotein transporter and possibly a zinc transporter. Ne's genome contain the most ABC transporters, including Iron(III), putracine, General L-amino acid, branched-chain amino acid, phosphate, lipoprotein, lipopolysaccharide, and possibly a molybdate transporter.

| | *Nephromyces* Alphaproteobacterial genome (Nαe) | *Nephromyces* Bacteroidetes Genome (NBe) | *Nephromyces* Betaproteobacterial Genome* (Nβe) |
|---|---|---|---|
| contigs | 11 | 1 | 1 |
| size in BP | 995,540 | 494,352 | 866,396 |
| CDS | 844 | 503 | 880 |
| tRNA | 35 | 31 | 40 |
| rRNA | 4 | 4 | 4 |
| GC content | 25% | 22% | 30% |

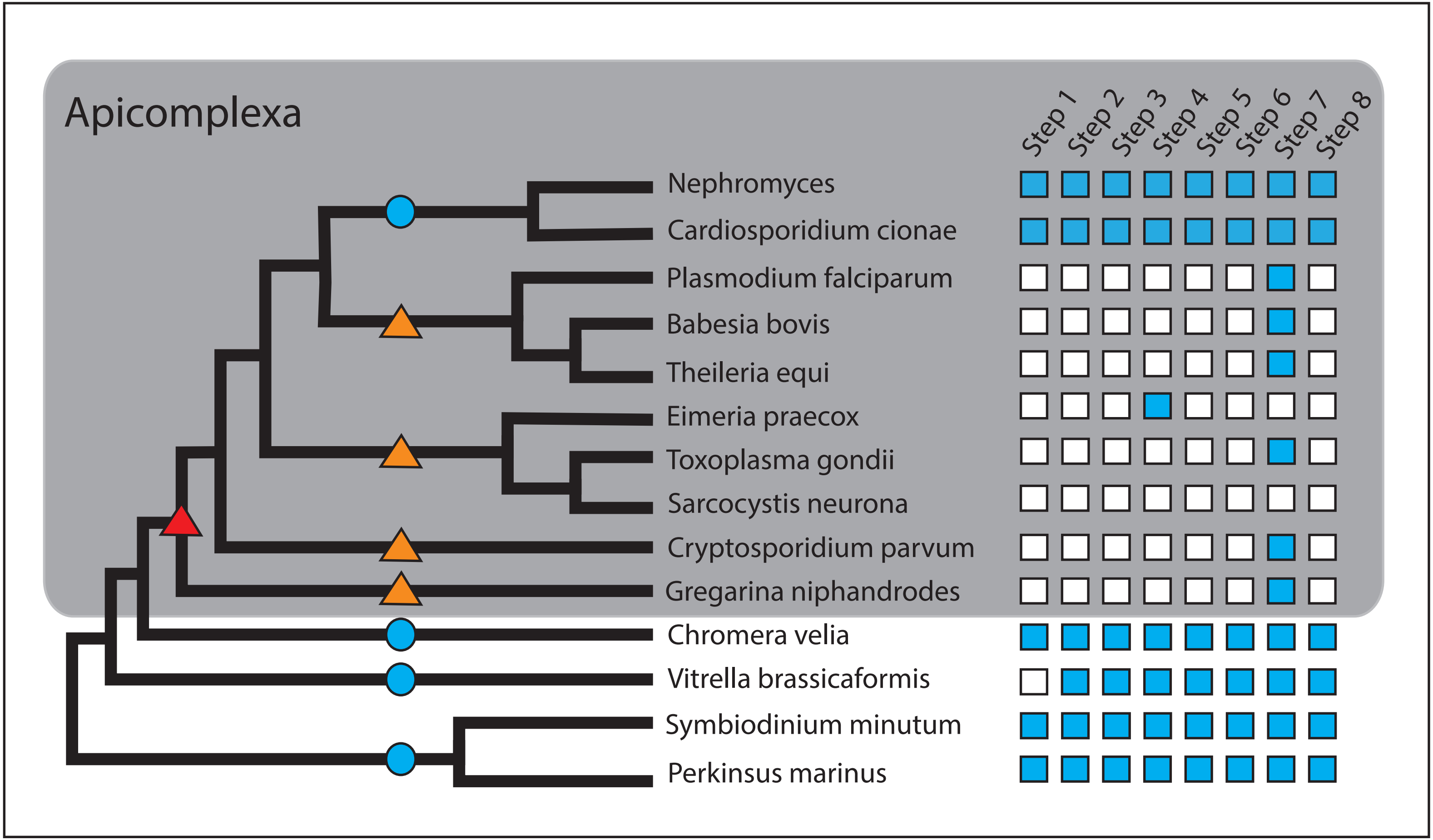
Apicomplexa
Step 1
Step 2
Step 3
Step 4
Step 5
Step 6
Step 7
Step 8
Nephromyces
Cardiosporidium cionae
Plasmodium falciparum
Babesia bovis
Theileria equi
Eimeria praecox
Toxoplasma gondii
Sarcocystis neurona
Cryptosporidium parvum
Gregarina niphandrodes
Chromera velia
Vitrella brassicaformis
Symbiodinium minutum
Perkinsus marinus

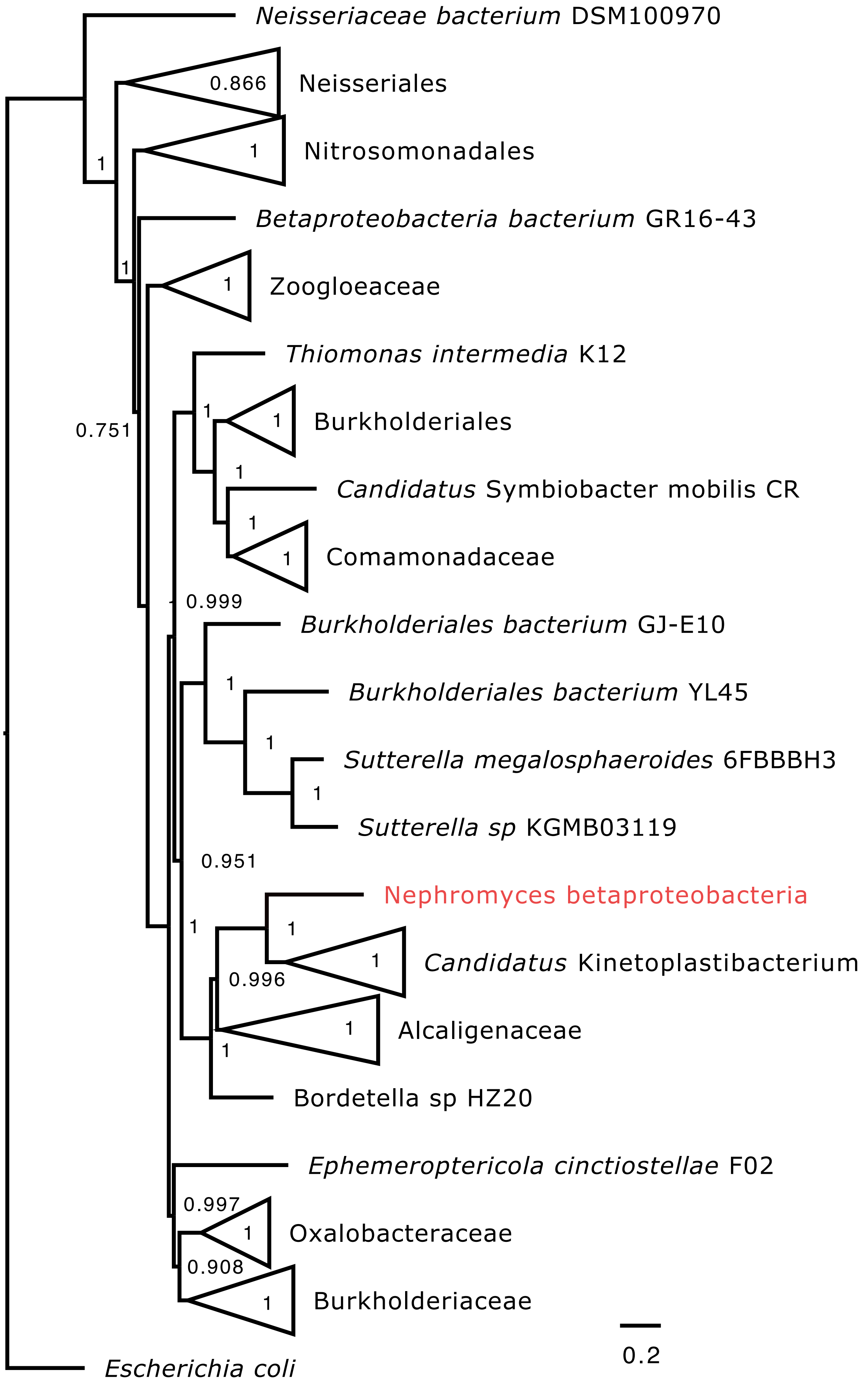

Neisseriaceae bacterium DSM100970
0.866
Neisseriales
1
Nitrosomonadales
1
Betaproteobacteria bacterium GR16-43
1
Zoogloeaceae
1
Thiomonas intermedia K12
1
Burkholderiales
1
0.751
1
Candidatus Symbiobacter mobilis CR
1
Comamonadaceae
1
0.999
Burkholderiales bacterium GJ-E10
1
Burkholderiales bacterium YL45
1
Sutterella megalosphaeroides 6FBBBH3
1
Sutterella sp KGMB03119
0.951
Nephromyces betaproteobacteria
1
1
Candidatus Kinetoplastibacterium
1
0.996
Alcaligenaceae
1
1
Bordetella sp HZ20
Ephemeroptericola cinctiostellae F02
0.997
Oxalobacteraceae
1
0.908
Burkholderiaceae
1
0.2
Escherichia coli

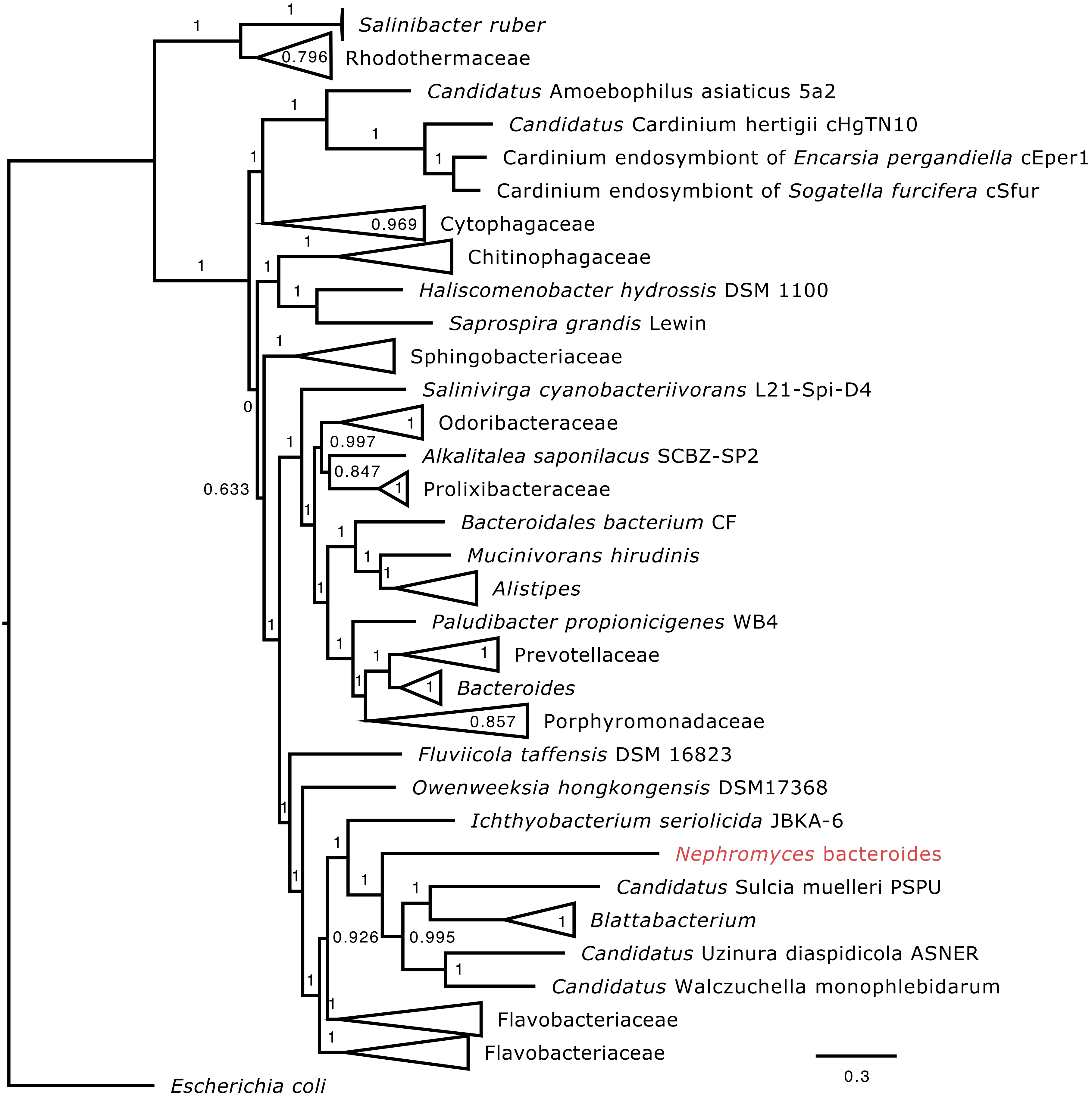
Salinibacter ruber
Rhodothermaceae
Candidatus Amoebophilus asiaticus 5a2
Candidatus Cardinium hertigii cHgTN10
Cardinium endosymbiont of Encarsia pergandiella cEper1
Cardinium endosymbiont of Sogatella furcifera cSfur
Cytophagaceae
Chitinophagaceae
Haliscomenobacter hydrossis DSM 1100
Saprospira grandis Lewin
Sphingobacteriaceae
Salinivirga cyanobacteriivorans L21-Spi-D4
Odoribacteraceae
Alkalitalea saponilacus SCBZ-SP2
Prolixibacteraceae
Bacteroidales bacterium CF
Mucinivorans hirudinis
Alistipes
Paludibacter propionicigenes WB4
Prevotellaceae
Bacteroides
Porphyromonadaceae
Fluviicola taffensis DSM 16823
Owenweeksia hongkongensis DSM17368
Ichthyobacterium seriolicida JBKA-6
Nephromyces bacteroides
Candidatus Sulcia muelleri PSPU
Blattabacterium
Candidatus Uzinura diaspidicola ASNER
Candidatus Walczuchella monophlebidarum
Flavobacteriaceae
Flavobacteriaceae
Escherichia coli
0.796
0.969
0.997
0.847
0.633
0
0.857
0.926
0.995
0.3

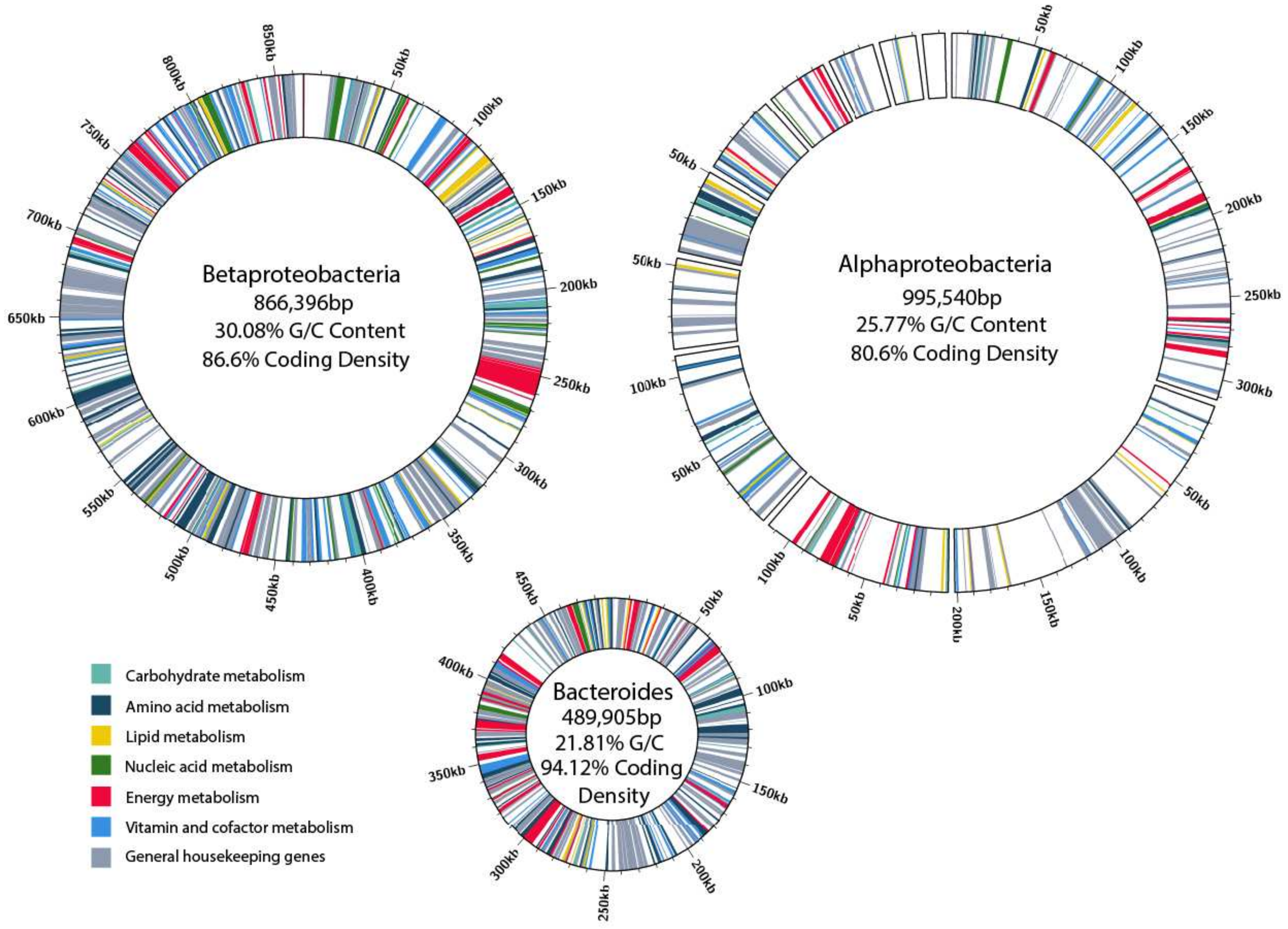
Betaproteobacteria
866,396bp
30.08% G/C Content
86.6% Coding Density
Alphaproteobacteria
995,540bp
25.77% G/C Content
80.6% Coding Density
Bacteroides
489,905bp
21.81% G/C
94.12% Coding
Density
Carbohydrate metabolism
Amino acid metabolism
Lipid metabolism
Nucleic acid metabolism
Energy metabolism
Vitamin and cofactor metabolism
General housekeeping genes

Supplementary Figure 1: De novo purine biosynthesis in Apicomplexans, Chromerids, and dinoflagellates. Genes are represented by boxes on the right side of the figure (blue=present, white=absent). The gray box encompasses Apicomplexa. Blue circles indicate complete de novo purine biosynthesis, the red triangle indicates presumed loss of purine synthesis in Apicomplexa, and the orange triangles indicate multiple independent losses of purine biosynthesis in Apicomplexa indicated by the presence of purine biosynthesis in Nephromycidae.

Supplementary Figure 2: Maximum likelihood tree of betaproteobacteria from GToTree pipeline with 1000 bootstrap values, including 203 genes and 471 genomes. *Escherichia coli* was used as an outgroup. Bootstrap support is listed on nodes.

Supplementary Figure 3: Maximum likelihood tree of Bacteroidetes from GToTree pipeline with 1000 bootstrap values, including 90 genes and 388 genomes. *Escherichia coli* was used as an outgroup. Bootstrap support is listed on nodes.

Supplementary Figure 4: Circos diagram of *Nephromyces'* bacterial endosymbionts showing overview of major functional categories (as predicted by KEGG), GC content, size, and coding density. Genomes are scaled relative to each other. The Alphaproteobacteria has not been assembled into a single molecule, and this figure also shows the size distribution of the 11 contigs making up this assembly.
